# The NKCC1 Antagonist Bumetanide Mitigates Interneuronopathy Associated with Ethanol Exposure *In Utero*

**DOI:** 10.1101/665851

**Authors:** Alexander G. J. Skorput, Stephanie M. Lee, Pamela W. L. Yeh, Hermes H. Yeh

**Affiliations:** Department of Molecular and Systems Biology, Geisel School of Medicine at Dartmouth, Hanover, NH 03755; Department of Neuroscience, School of Medicine, University of Minnesota Twin Cities, Minneapolis, MN 55455

**Author notes:** Corresponding Author: Hermes H. Yeh, Ph.D., Department of Molecular and Systems Biology, Geisel School of Medicine at Dartmouth, 66 College Street, Hanover, NH 03755. These authors contributed equally to this work (Alexander G. J. Skorput, Stephanie M. Lee).

## Abstract

Prenatal exposure to ethanol induces aberrant tangential migration of corticopetal GABAergic interneurons, and long-term alterations in the form and function of the prefrontal cortex. We have hypothesized that interneuronopathy contributes significantly to the pathoetiology of fetal alcohol spectrum disorders (FASD). Activity-dependent tangential migration of GABAergic cortical neurons is driven by depolarizing responses to ambient GABA present in the cortical enclave. We found that ethanol exposure potentiates the depolarizing action of GABA in GABAergic cortical interneurons of the embryonic mouse brain. Pharmacological antagonism of the cotransporter NKCC1 mitigated ethanol-induced potentiation of GABA depolarization and prevented aberrant patterns of tangential migration induced by ethanol *in vitro*. In a model of FASD, maternal bumetanide treatment prevented interneuronopathy in the prefrontal cortex of ethanol exposed offspring, including deficits in behavioral flexibility. These findings position interneuronopathy as a mechanism of FASD symptomatology, and posit NKCC1 as a pharmacological target for the management of FASD.

## Introduction

Binge-type exposure of the embryonic mouse brain to a moderate level of ethanol leads to interneuronopathy, hallmarked by aberrant tangential migration of corticopetal GABAergic interneurons derived from the medial ganglionic eminence (MGE) (Skorput et al., 2015). Interneuronopathies are widely implicated in neurodevelopmental disorders (Catterall, 2018; Kato and Dobyns, 2005; Katsarou et al., 2017), including Fetal Alcohol Spectrum Disorders (FASD) (Skorput et al., 2015; Skorput and Yeh, 2016). Data from the Behavioral Risk Factor Surveillance System for the years 2015 – 2017 show self-reported rates of current drinking, and binge drinking (≥ 4 drinks) within the past 30 days to be 11.5% and 3.9%, respectively, among pregnant woman aged 18 - 44 years. Pregnant binge drinkers averaged 4.5 binge episodes in the past 30 days with an average binge intensity of 6.0 drinks (Denny et al., 2019). As a result of such binge drinking episodes, FASD remains a major public health concern in the United States, and a recent assessment of first-graders in 4 US communities conservatively estimated the prevalence rates of FASD to range from 1.1% to 5.0% (May et al., 2018). The global prevalence rate of FASD is estimated at 7.7 per 1000 children, and is as high as 111.1 per 1000 children among some populations (Lange et al., 2017). Despite this prevalence, our understanding of the embryonic pathoetiology of FASD is incomplete, hampering development of targeted medication and broadly applicable treatment strategies.

The more efficacious current treatments for FASD rely primarily on postnatal behavioral therapy (Nash et al., 2017), or pharmaceutics aimed to treat the symptoms, rather than the etiology, of this neurodevelopmental disorder (Peadon et al., 2009; Rowles and Findling, 2010). Indeed, there is a societal need to uncover the cellular and subcellular underpinnings by which ethanol adversely affects neurodevelopmental processes (Ismail et al., 2010; Kodituwakku, 2010; Pruett et al., 2013). Here we report on pharmacologic mitigation of ethanol-induced interneuronopathy and associated executive function deficits in a preclinical model of FASD.

GABAergic neuroblasts arising in the MGE are fated to become parvalbumin (PV^+^) or somatostatin expressing inhibitory interneurons in the cortex. Approximately 50% of this MGE-derived population will mature to become PV^+^ fast-spiking basket cells which are electrically coupled across the deep layers of the neocortex and pace oscillatory gamma rhythms, which are required for higher cognition (Batista-Brito and Fishell, 2009; Xu et al., 2008). Thus, the corticopetal tangential migration and maturation of MGE-derived GABAergic interneurons is critical to the establishment of inhibitory/excitatory balance in intracortical circuits, and proper cortical functioning (Batista-Brito and Fishell, 2009; Catterall, 2018).

Building on our previous work (Cuzon et al., 2008, 2006), we hypothesized that ethanol’s positive allosteric modulation of GABA_A_ receptors (GABA_A_R) expressed on migrating neuroblasts is the mechanism by which *in utero* ethanol exposure causes enduring interneuronopathy in models of FASD (Skorput et al., 2015; Skorput and Yeh, 2016). In these immature neurons, GABA_A_R activation results in membrane depolarization (Ben-Ari, 2014, 2002; Owens and Kriegstein, 2002). This paradoxical depolarizing action has been postulated to drive the corticopetal tangential migration of GABAergic interneurons during corticogenesis (Behar et al., 1996; Ben-Ari, 2002; Ben-Ari et al., 2012; Cuzon et al., 2006; Wang and Kriegstein, 2009). To test the influence of GABAergic depolarization on ethanol induced interneuronopathy, we sought to normalize ethanol’s potentiation of GABAergic depolarization by reducing the driving force of GABA-induced membrane depolarization via a decrease in the intracellular concentration of the GABA_A_R permeable anion Cl^-^ ([Cl^-^]_i_).

To this end, we targeted the chloride importing Na^+^-K^+^-2Cl^-^ isoform 1 cotransporter (NKCC1) because it is expressed at high levels in the embryonic brain when its activity predominates over that of the chloride exporting K^+^-Cl^-^ co-transporter (KCC2). This differential activity results in a net higher [Cl^-^]_i_ compared to [Cl^-^]_o_. As such, the reversal potential for chloride, and that of GABA-activated responses (E_GABA_), is set at a membrane potential that is depolarized relative to the resting membrane potential. Thus, GABA_A_R activation causes chloride extrusion and membrane depolarization in these migrating neuroblasts (Ben-Ari, 2014, 2002; Owens and Kriegstein, 2002). The loop diuretic bumetanide is an NKCC1 antagonist that shifts the E_GABA_ of embryonic neuroblasts to a more negative membrane potential when administered maternally (Wang and Kriegstein, 2011). Given these considerations, we tested the hypothesis that *in utero* treatment with bumetanide during the period of prenatal ethanol exposure will mitigate manifestation of prenatal ethanol-induced aberrant tangential migration, and prevent deficits in behavioral flexibility.

The timing of gestational exposure to ethanol is a key determinant of the diagnostic outcome of FASD (May et al., 2013; Pruett et al., 2013). Corticogenesis occurs in earnest during the mid-first trimester when the developing brain is highly vulnerable to insult by ethanol (Ayoola et al., 2009; Clancy et al., 2001; May et al., 2014). We established a mouse model that simulates an early gestational, mid-first trimester human equivalent, exposure to binge-type maternal ethanol consumption from embryonic day (E) 13.5 - E16.5 (Clancy et al., 2001; Skorput et al., 2015). Using this model we demonstrated enhanced entry of MGE-derived GABAergic interneurons into the prefrontal cortex (PFC), a persistent increase in the number of PV^+^ interneurons in the young adult PFC, and impairment in the PFC-dependent behavioral flexibility of offspring (Skorput et al., 2015).

In the present study, we used this mouse model to assess whether the NKCC1 cotransporter is a tractable pharmacological target for normalizing *in utero* ethanol exposure-induced escalation of tangential migration to the prefrontal cortex. We report here that ethanol induces a shift in E_GABA_ toward a more depolarized potential, accounting at least in part for ethanol’s potentiation of GABA-induced depolarizing responses in embryonic MGE-derived GABAergic interneurons. In the short term, antagonizing NKCC1 with bumetanide mitigated ethanol-induced aberrant tangential migration. In the long-term, bumetanide treatment *in vivo* prevented prenatal ethanol exposure-induced interneuronopathy in the prefrontal cortex, and the associated deficits in PFC-dependent behavioral flexibility. Our findings support the feasibility of a pharmacological strategy to target NKCC1 for the management of FASD.

## Results

### Ethanol induces a depolarizing shift in the GABA reversal potential of embryonic MGE-derived GABAergic cortical interneurons that is normalized by the NKCC1 inhibitor bumetanide

Nkx2.1^+^ embryonic MGE-derived GABAergic cortical interneurons were identified by tdTomato fluorescence and targeted for perforated patch clamp recording in acute (200 μm) telencephalic slices obtained from E14.5 - 16.5 Nkx2.1Cre/Ai14 mouse brain (Fig. 1a). GABA (50 μM), ethanol (6.5 mM) and aCSF were loaded into separate barrels of a multi-barrel drug pipette and focally applied by regulated pressure either individually, or in combination, in the immediate vicinity of the cell under study (Fig. 1a). Since GABA-induced membrane depolarization is prevalent in many types of immature neurons early in brain development, and has been postulated to promote the tangential migration of GABAergic cortical interneurons (Behar et al., 1996; Ben-Ari, 2002; Ben-Ari et al., 2012; Cuzon et al., 2006; Wang and Kriegstein, 2009), we asked whether exogenously-applied GABA depolarized embryonic Nkx2.1^+^ cortical interneurons. In Figure 1b, the representative sets of digitized raw traces display current responses to 500 ms pressure applications of 50 μM GABA monitored at varying holding potentials before (black traces) and during ethanol exposure (red traces). The polarity and peak amplitude of each GABA response were plotted as a function of the corresponding holding potential, and the intercept of the current-voltage plot along the abscissa was used to estimate E_GABA_. Figure 1c shows linear regression of the group mean data. The control mean E_GABA_ (dotted black line in Fig. 1c) was -23.18 ± 1.45 mV, which was significantly more depolarized than the mean resting membrane potential, estimated in whole-cell mode (dotted blue line in Fig. 1c; -41.9 ± 2.3 mV; unpaired t-test, P<0.01). Importantly, Figure 1c illustrates that ethanol exposure shifted E_GABA_ rightward relative to that assessed during the control epoch (dotted red line). Figure 1d illustrates the mean change in E_GABA_ for each litter before and during ethanol exposure, indicating that the mean E_GABA_ of control cells was shifted to significantly more depolarized membrane potentials with exposure to 6.5 mM ethanol (Fig. 1d; -16.52 ± 0.93 mV; P=0.0043). The finding that ethanol exposure results in a depolarizing shift in E_GABA_ suggests that it increases [Cl^-^]_i_ and thus, chloride drive.

**Figure 1.**
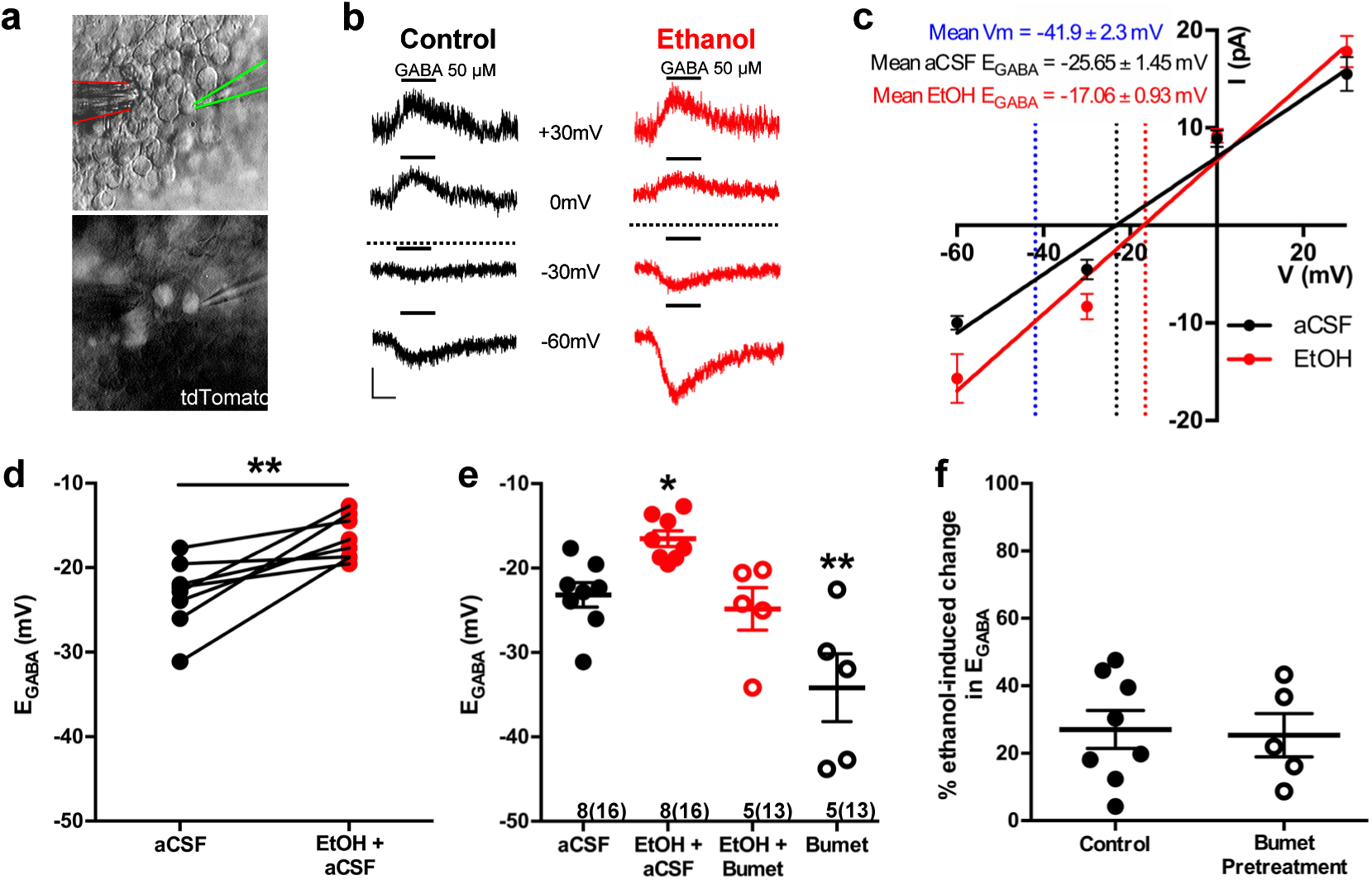
Ethanol induces a depolarizing shift in GABA reversal potential in embryonic MGE-derived GABAergic cortical interneurons that is normalized by the NKCC1 inhibitor bumetanide. a) Hoffman modulated contrast image of acute telencephalic slice of E14.5 mouse brain showing patch clamp electrode (green outline) holding a tdTomato fluorescent Nkx2.1^+^ MGE-derived GABAergic interneuron in the perforated-patch voltage clamp configuration, with a multi-barrel drug pipette (red outline) positioned in the vicinity. b) E_GABA_ was empirically determined by focal application of GABA (black bars, 50 µM) at varying holding potentials under control (black traces), and ethanol exposure (6.5mM EtOH, red traces), conditions. Dotted lines indicate E_GABA_ for each condition, scale bar vertical = 10 pA; horizontal = 500 ms. c) I/V plot of peak GABA induced current over holding potential defines E_GABA_ as the x-intercept under control (black) and ethanol exposure (red) conditions. Dotted blue line denotes mean resting membrane potential. d) E_GABA_ of neurons under control conditions (aCSF) and during ethanol exposure (EtOH+aCSF). ** = P<0.01 paired t-test. e) Group-wise comparison of E_GABA_ in Nkx2.1^+^ neurons without and with bumetanide pretreatment (Bumet, 20 µM) and during exposure to EtOH with bumetanide pretreatment (EtOH+Bumet). Numbers below scatter denote number of litters and (cells recorded). *, ** = P<0.05, P<0.01 compared to aCSF, one-way ANOVA with Dunnett post-test. f) Magnitude of change in E_GABA_ induced by 6.5mM ethanol without and with bumetanide pretreatment (P>0.05, unpaired t-test).

Pretreatment of acute E14.5 - 16.5 telencephalic slices for 30 minutes with the NKCC1 antagonist bumetanide (20 µM) normalized the ethanol-induced depolarization of E_GABA_ to control levels (Fig. 1e; EtOH+Bumet; -24.82 ± 2.52 mV; n=5 litters, 13 cells total; P = 0.56). The basis of this normalization is likely a functional antagonism as bumetanide pretreatment significantly hyperpolarized E_GABA_ in Nkx2.1^+^ GABAergic interneurons compared to controls (Fig. 1e; Bumet; -34.17 ± 4.02 mV; n=5 litters, 13 cells total; P = 0.004), and the magnitude of the depolarizing shift in E_GABA_ induced by exposure to 6.5mM ethanol was not effected by bumetanide pretreatment (Fig. 1f; Control; 27.08 ± 5.61 mV; Bumet pretreatment; -25.38 ± 6.41 mV; P=0.37). These results prompted us to ask whether bumetanide treatment may restrict the enhanced chloride driving force caused by ethanol-induced increases in [Cl^-^]_i_ to thus mitigate ethanol’s potentiation of depolarizing GABA_A_R activity in embryonic GABAergic interneurons.

### Bumetanide attenuates ethanol-induced potentiation of depolarizing GABA responses in embryonic MGE-derived GABAergic cortical interneurons

In perforated patch-clamp recordings, we analyzed current response amplitudes of GABAergic cortical interneurons held at -60 mV to 500 ms applications of 50 μM GABA before and during 6.5 mM ethanol exposure in E14.5-E16.5 telencephalic slices (Fig. 2a, inset). Figure 2a illustrates that, relative to baseline conditions, the mean GABA response amplitude during ethanol exposure is significantly greater (aCSF; 10.02 ± 0.99 pA; EtOH+aCSF; 16.89 ± 2.39 pA; n = 8 litters, 17 cells total; P = 0.0021). The mean amplitude of the GABA-activated current responses of GABAergic interneurons in the bumetanide pretreated slices was 10.53 ± 0.62 pA before ethanol exposure, and was significantly increased to 13.45 ± 1.31 pA during ethanol exposure (Fig. 2b; n=5 litters, recorded from 11 cells; P = 0.019). Overall, ethanol exposure potentiated the amplitude of GABA responses by 59.37 ± 8.83% under control conditions, and by 26.46 ± 6.26% in the bumetanide pretreatment group, demonstrating the ability of bumetanide to significantly decrease the mean ethanol-induced change in GABA response (Fig. 2c; P=0.011). Taken together with our earlier finding that GABA-activated current responses in embryonic MGE-derived GABAergic interneurons are mediated by GABA_A_ receptors (Cuzon Carlson and Yeh, 2011; Cuzon et al., 2008), these results indicate that NKCC1 antagonism attenuates ethanol-induced potentiation of depolarizing GABA_A_R-activated responses in migrating GABAergic cortical interneurons.

**Figure 2.**
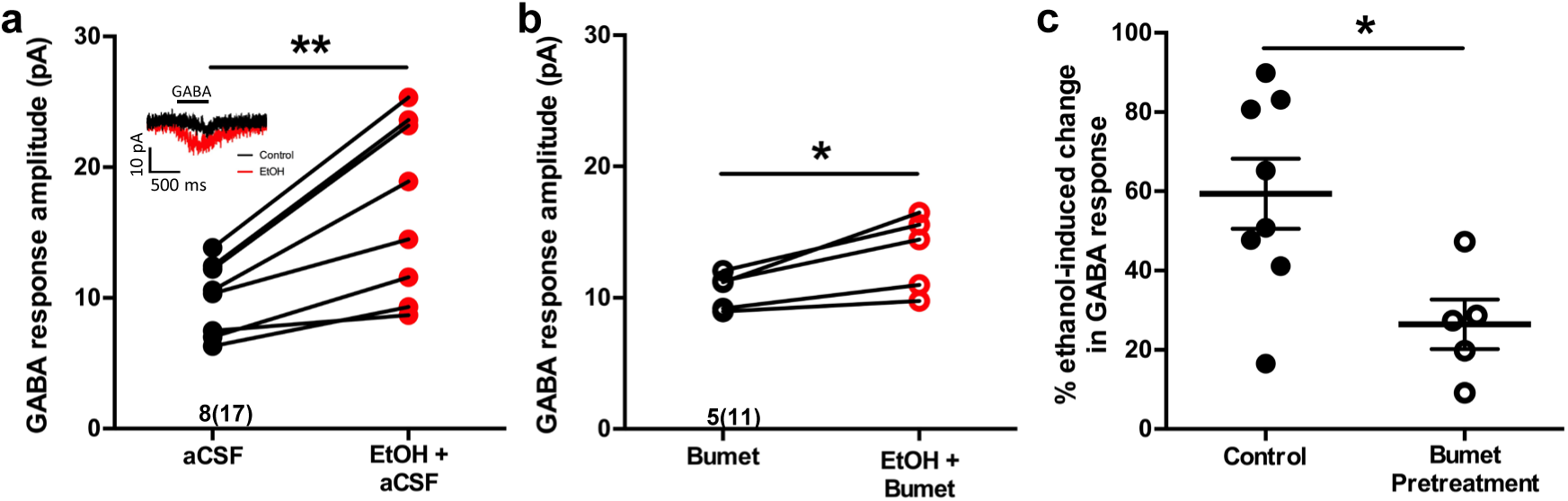
Bumetanide attenuates ethanol-induced potentiation of depolarizing GABA responses in embryonic MGE-derived GABAergic cortical interneurons. **a)** Peak current amplitude recorded from Nkx2.1^+^ MGE-derived GABAergic interneurons in slices of E14.5 mouse telencephalon in response to focal application of GABA (50 µM) under control (aCSF; inset: black trace) or ethanol exposure (EtOH, 6.5 mM; inset: red trace), scale bar vertical = 10 pA; horizontal = 500 ms. ** = P<0.01 paired t-test. **b)** Peak current amplitude in response to focal application of GABA (50 µM) in the presence of bumetanide (Bumet) or ethanol and bumetanide (EtOH+Bumet). * = P<0.01 paired t-test. **c)** Percent change in peak GABA response induced by ethanol exposure at baseline (aCSF) and in the presence of bumetanide (Bumet). * = P <0.05 unpaired t-test.

### Co-treatment with bumetanide prevents ethanol’s enhancement of tangential migration in vitro

We then asked whether NKCC1 antagonism inhibits ethanol-induced escalation of tangential migration among cortical GABAergic interneurons (Cuzon et al., 2008; Skorput et al., 2015; Skorput and Yeh, 2016). To this end, organotypic slice cultures containing the embryonic PFC were prepared from E14.5 Nkx2.1Cre/Ai14 brains, incubated in control or ethanol-containing medium, and tangential migration of MGE-derived interneurons was assessed without or with co-exposure to bumetanide (Fig. 3). Nkx2.1^+^ GABAergic interneurons were counted in consecutive bins 100 µm in width arranged ventro-dorsally and beginning at the cortico-striate junction (Fig. 3a). Relative to controls, organotypic telencephalic slices cultured in media containing 50 mM ethanol had more Nkx2.1^+^ GABAergic interneurons per 100 μm cortical bin in the embryonic PFC (Fig. 3a&b; Control 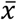= 61.3 ± 2.8 cells, 12 cultures; EtOH 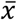= 78.5 ± 3.5 cells, 10 cultures; P = 0.00166). The addition of bumetanide (20μM) to the culture medium prevented the ethanol-induced escalation of Nkx2.1^+^ interneuron migration into the cortex with no significant difference observed compared to control (Fig. 3a&b; EtOH+Bumet 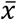= 61.1 ± 2.4 cells, 10 cultures; P >0.999). Bumetanide alone (20 µM) did not affect Nkx2.1^+^ interneuron entry into the cortex compared to controls (Fig. 3a&b; Bumet 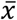= 60.3 ± 3.7 cells, 9 cultures; P >0.999). When analyzed in terms of the number of Nkx2.1^+^ cells per cortical bin, the largest increase in Nkx2.1^+^ cell number was in the cortical region most proximal to the corticostriate juncture (Fig. 3c). These results are consistent with aberrant tangential migration being a primary effect of ethanol exposure in the fetal brain. In addition, these data demonstrate that bumetanide can inhibit the ethanol-induced supranormal corticopetal migration of GABAergic interneurons at concentrations that did not affect their normal pattern of migration.

**Figure 3.**
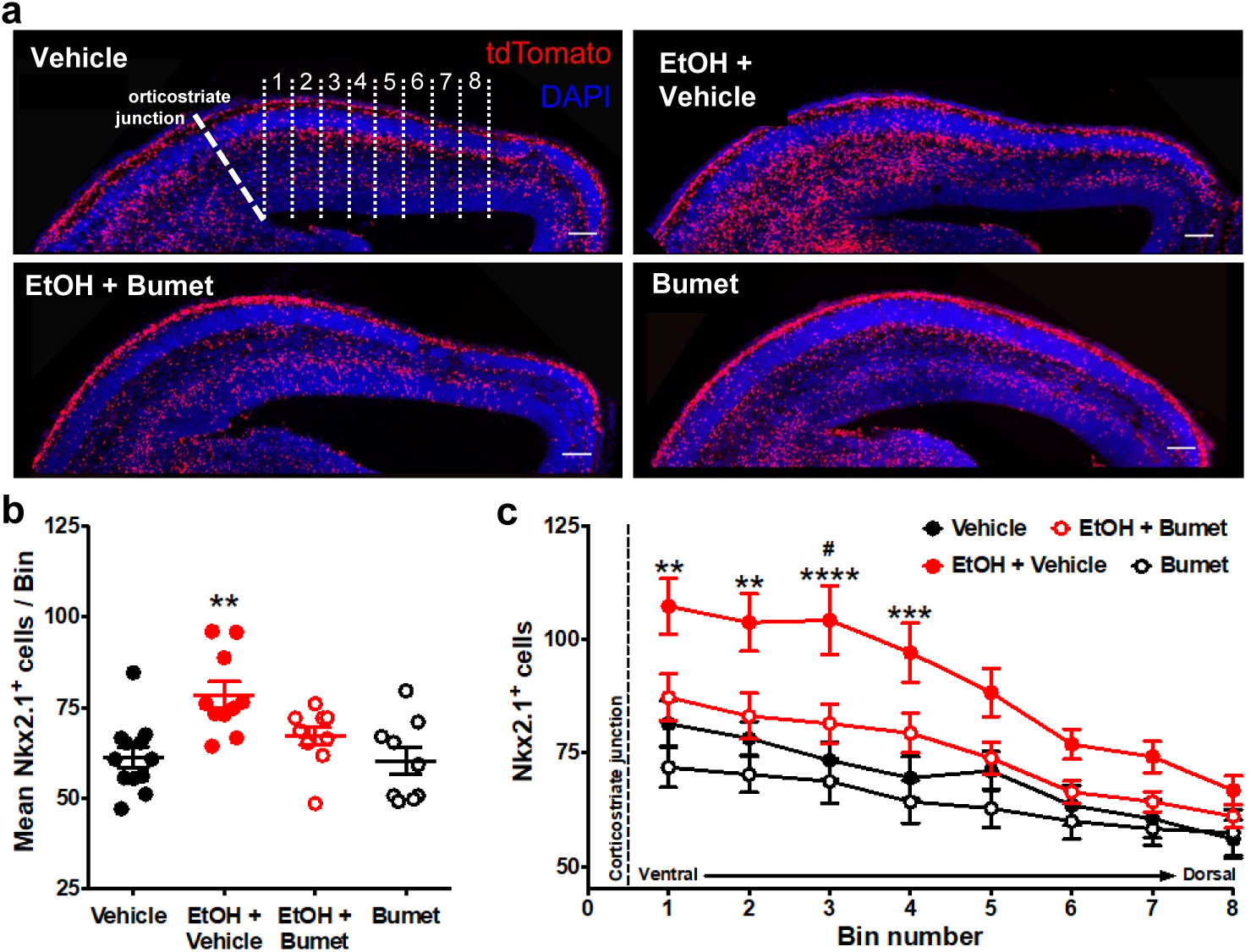
Co-treatment with bumetanide prevents ethanol’s enhancement of tangential migration in vitro. **a)** Fluorescent images of organotypic E14.5 Nkx2.1-Cre/Ai14 mouse telencephalic brain slices treated with vehicle, ethanol (50 mM) + vehicle (EtOH + Vehicle), ethanol + bumetanide (EtOH+Bumet, 20 µM), or bumetanide alone (Bumet, 20 µM), and counterstained with DAPI (blue). The corticostriate junction is marked by the dashed line, and numbered counting bins are denoted by dotted lines in the vehicle panel. Scale bars = 100 µm. **b)** Mean number of tdTomato flourecent Nkx2.1^+^ cells per 100 µm bin. ** = P<0.01 one-way ANOVA with Bonferroni post-test. Each dot represents one organotypic culture. **c)** Number of tdTomato fluorescent Nkx2.1^+^ cells in each 100 µm bin. **, ***, **** = P<0.01, 0.001, 0.0001 compared to control, # = P<0.05 compared to EtOH+Bumet two-way ANOVA with Bonferroni post-test.

### Maternal bumetanide treatment prevents ethanol-induced escalation of tangential migration in vivo

To assess the short term interaction between binge ethanol exposure and bumetanide treatment *in vivo*, we co-treated binge-type ethanol-consuming (5% EtOH w/w) pregnant dams harboring Nkx2.1Cre/Ai14 embryos daily with bumetanide (i.p.; 0.15mg/kg dissolved in DMSO) from E13.5 – 16.5 and analyzed the density of Nkx2.1^+^ profiles in the embryonic PFC at E16.5 (Fig. 4a). Ethanol exposure alone significantly increased the number of Nxk2.1^+^ neurons in the dorsomedial telencephalon (Fig. 4b; Control 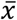= 1.78×10^-3^ ± 5.90×10^-5^ cells/μm^2^, 10 brains from 3 litters; EtOH 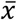= 2.61×10^-3^ ± 1.39×10^-4^ cells/μm^2^, 10 brains from 4 litters; P = 1.13×10^-5^). Maternal treatment over the course of binge-type ethanol exposure with bumetanide significantly attenuated the ethanol-induced increase in MGE-derived GABAergic interneuron density in the embryonic PFC (Fig. 4b; EtOH+Bumet DMSO i.p. 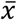= 1.99×10^-3^ ± 8.79×10^-5^ cells/μm^2^, 8 brains from 3 litters; P = 0.00357). In addition, bumetanide co-treatment completely normalized the ethanol-induced effect, returning MGE-derived interneuron density to control levels (Fig. 4b; P >0.999). As a negative control, given that bumetanide is relatively insoluble in aqueous solutions, treatment with normal saline to which bumetanide (0.015 mg/ml) was added failed to decrease the ethanol induced enhancement of tangential migration when injected i.p. (Fig. 4b; EtOH+Bumet NS 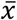= 2.49×10^-3^ ± 1.45×10^-5^ cells/μm^2^, 5 brains from 2 litters; P >0.999), and a significant increase in Nkx2.1^+^ cell density persisted in the E16.5 PFC compared to controls (Fig. 4b; P = 0.00481).

**Figure 4.**
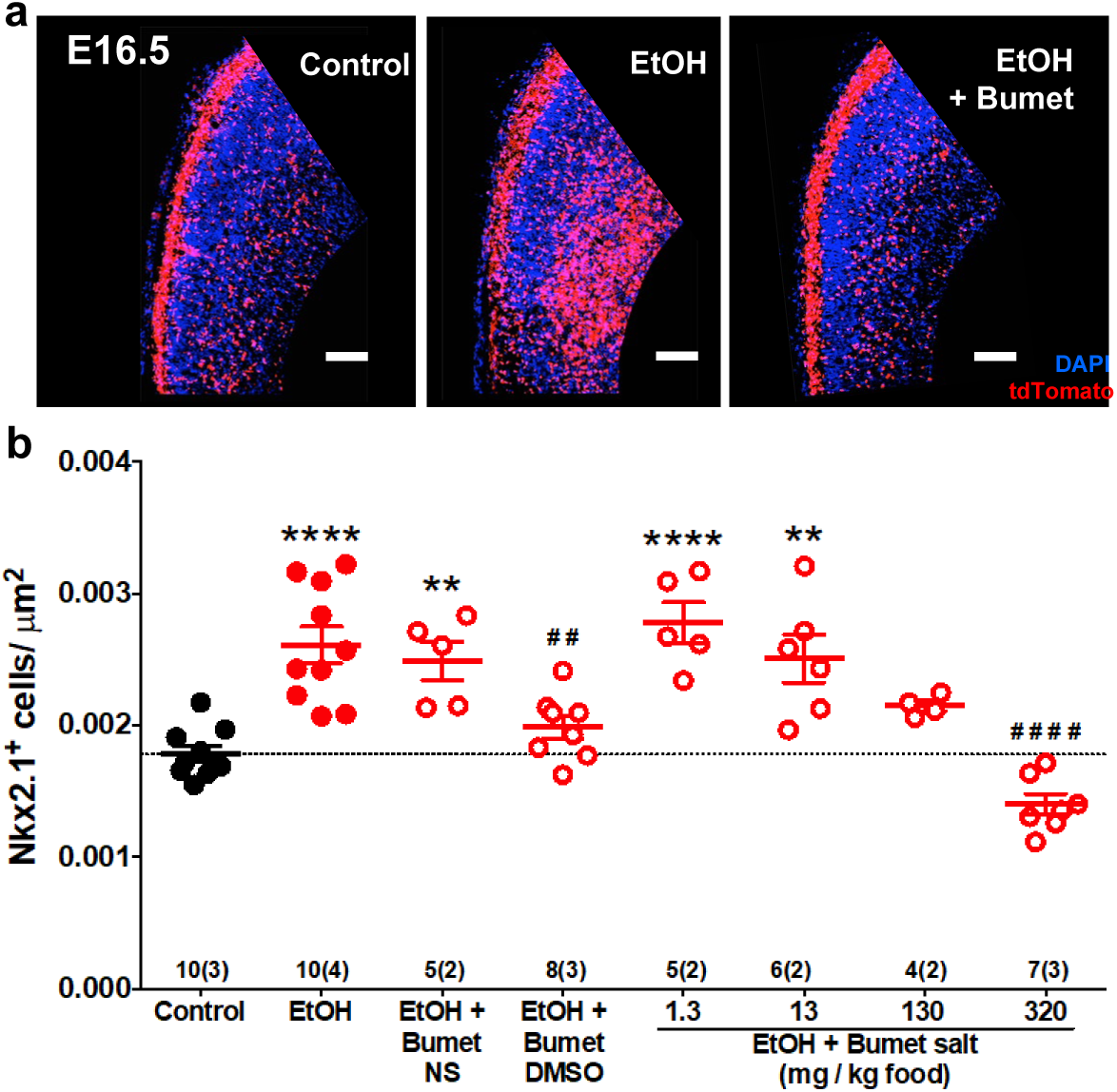
Maternal bumetanide treatment prevents ethanol-induced escalation of tangential migration in vivo. Nkx2.1^+^ cells were identified by tdTomato fluorescence and quantified in the dorso-medial telencephalon of E16.5 embryonic brain. **a)** fluorescent images counterstained with DAPI following control, binge-type maternal ethanol consumption from E13.3 - E16.5 (EtOH), or ethanol exposure in combination with maternal bumetanide treatment administered i.p. 0.15 mg/kg/day (E13.5 - E16.5) dissolved in DMSO (EtOH+Bumet). Scale bars = 100 µm. **b)** Quantification of Nkx2.1^+^ cells in the dorso-medial telencephalon after ethanol exposure in combination with maternal bumetanide treatment administered i.p. 0.15 mg/kg/day (E13.5 - E16.5) dissolved in normal saline (EtOH+Bumet NS) or DMSO (EtOH_Bumet DMSO), or maternal bumetanide treatment administered in the ethanol containing liquid diet (EtOH+Bumet salt; 1.3, 13, 130, 320 mg/kg food), or control conditions under which dams consumed equicaloric liquid food from E13.5-E16.5 (control). Dotted line highlights mean of control group. Numbers above x-axis denote number of brains and (number of litters). **,**** = P<0.01, 0.0001 compared to control; ##, #### = P<0.01, 0.0001 compared to EtOH; one-way ANOVA with Bonferroni post-test.

We next sought to simulate a more clinically relevant route of bumetanide administration. We asked whether bumetanide, administered orally in the course of *ad lib* feeding, can prevent the *in vivo* effects of binge-type ethanol exposure on the entry of MGE-derived interneurons to the embryonic PFC. To this end, a Na^+^ bumetanide salt (320, 130, 13, 1.3 mg/kg food) was dissolved in the 5% w/w ethanol containing liquid diet fed to binge ethanol-consuming dams.

E16.5 embryos from dams ingesting ethanol-containing liquid food with 320 mg bumetanide salt /kg food had a significantly lower density of Nkx2.1^+^ cells in the PFC compared to ethanol exposure alone (Fig 4b; EtOH+Bumet 320mg/kg food = 1.40×10^-3^ ± 7.96×10^-5^ cells/μm^2^, 7 brains from 2 litters; P = 1.38×10^-8^) with complete normalization relative to the control cohort (Fig 4b; P = 0.483). At a concentration of 130 mg bumetanide salt /kg food, no significant difference in Nkx2.1^+^ cell density was found in the E16.5 PFC compared to control or binge ethanol exposure alone (Fig. 4; EtOH+Bumet 130mg/kg food = 2.15×10^-3^ ± 4.07×10^-5^ cells/μm^2^, 4 brains from 2 litters; P > 0.999, P = 0.465 respectively). Embryos born to dams that consumed a liquid diet containing both 5% w/w ethanol and the lowest concentrations of bumetanide tested (1.3 or 13mg bumetanide salt /kg food) showed significantly higher densities of MGE-derived GABAergic interneurons in the PFC compared to controls (Fig. 4b; EtOH+Bumet 1.3mg/kg food 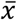= 2.78×10^-3^ ± 1.55±10^-4^ μm^2^, 5 brains from 2 litters; EtOH+Bumet 13mg/kg food 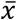= 2.50×10^-3^ ± 1.81×10^-4^ cells/μm^2^, 6 brains from 2 litters; P = 1.65×10^-5^, P = 0.00152 respectively) and Nkx2.1^+^ cell densities were not significantly decreased compared to EtOH exposure alone (Fig 4b; P >0.999). Thus, maternal bumetanide treatment dose dependently mitigates binge-type ethanol-induced aberrant migration of embryonic MGE-derived interneurons in the embryonic brain.

### Treatment of binge ethanol consuming dams with bumetanide prevents interneuronopathy in the PFC of young adult offspring

To determine the long-term effect of NKCC1 antagonism on ethanol-induced interneuronopathy, we quantified the distribution of PV^+^ interneurons in the PFC of young adult mice that had been exposed prenatally to binge-type ethanol without or with bumetanide co-treatment (daily maternal i.p. injection, 0.15mg/kg, E13.5 - E16.5). The PFC was divided into subregions (anterior cingulate cortex (ACC), prelimbic (PL), infralimbic (IL)), and subdivided by cortical layer, based on the cytoarchitecture as revealed by DAPI counterstaining. Previously, we reported a layer V-specific increase in PV^+^ interneurons in the young adult mPFC following binge-type ethanol exposure *in utero* (Skorput et al., 2015). In comparison, the number of PV^+^ interneurons in the PFC of binge-type ethanol-exposed offspring treated *in utero* with bumetanide was significantly lower in layer V of the ACC (Fig. 5a&b; EtOH 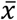= 47.7 ± 3.5, 9 brains from 4 litters; EtOH+Bumet 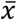= 37.3 ± 2.0, 4 brains from 2 litters; P = 0.00429) and the PL region (Fig. 5a&c; EtOH+Bumet 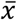= 30.9 ± 3.0, 9 brains from 4 litters; EtOH+Bumet 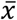= 19.1 ± 1.5, 4 brains from 2 litters; P = 3.27×10^-5^). The increase in PV^+^ interneurons of the ACC induced by *in utero* ethanol exposure was completely prevented by bumetanide co-treatment; there was no significant difference in the number of PV^+^ interneurons in layer V of the ACC between offspring exposed to ethanol *in utero* that received maternal bumetanide treatment and controls (Fig. 5a&b; Control 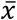= 31.8 ± 2.0, 11 brains from 4 litters; P = 0.384). Maternal bumetanide treatment also completely prevented the ethanol induced effect in layer V of the PL region (Fig. 5a&c; Control 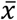= 19.7 ± 1.9, 11 brains from 4 litters; P >0.999). The effect of bumetanide treatment on preventing *in utero* ethanol-induced interneuronopathy extended throughout layer V of the PFC, with no significant difference in the number of PV^+^ interneurons in the IL region compared to controls (Fig. 5a&d; Control 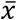= 8.4 ± 0.7, 11 brains from 4 litters; EtOH+Bumet 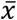= 10.4 ± 2.1, 4 brains from 2 litters; P = 0.228). Additionally, maternal treatment with bumetanide (0.15 mg/kg i.p. daily, E14.5 - E16.5) in the absence of *in utero* binge-type ethanol exposure (control liquid diet) did not alter the number of PV^+^ interneurons in any of the PFC regional layers analyzed in young adult offspring (Fig. 5a-d; Bumet, 3 brains from 2 litters; P >0.247).

**Figure 5.**
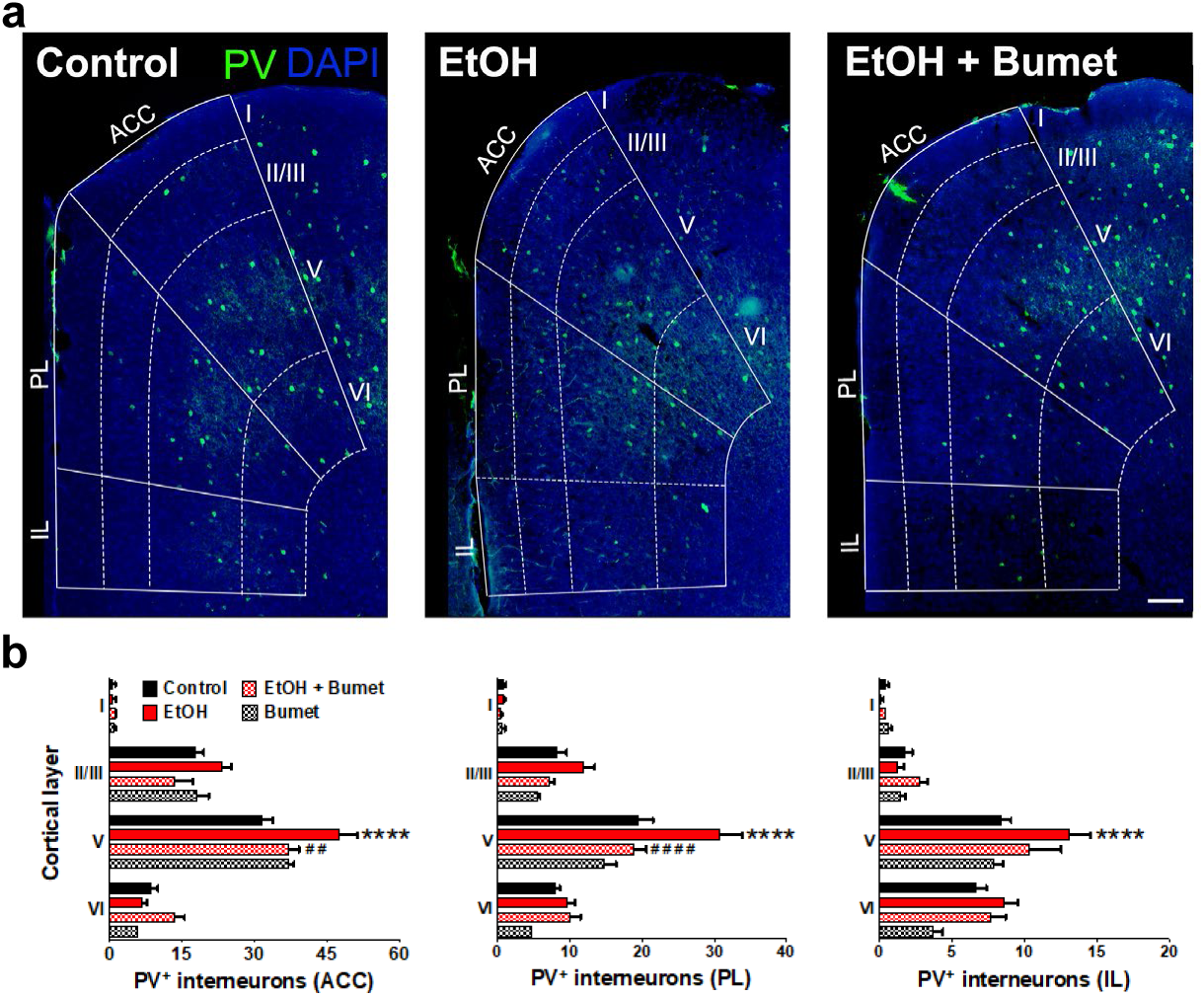
Treatment of binge ethanol-exposed dams with bumetanide prevents the interneuronopathy associated with ethanol exposure *in utero*. **a)** Histological sections of young adult mouse prefrontal cortex processed for parvalbumin immunofluorescence, counterstained with DAPI, and binned by functional region and layer using DAPI cytoarchitecture. Scale bar = 100 µm. **b)** Quantification of PV^+^ MGE-derived GABAergic cortical interneurons in mice without (control), and with (EtOH) binge-type ethanol exposure (E13.5 - E16.5), as well as mice with ethanol exposure born to dams treated with bumetanide (0.15 mg/kg/day i.p. in DMSO; EtOH+Bumet), or mice born to dams that consumed control diet and received bumetanide treatment (Bumet). **** = P<0.0001 compared to control; ##, #### = P<0.01, 0.0001 compared to EtOH; two-way ANOVA with Bonferroni post-test analyzed per region, and stratified by treatment and cortical layer. ACC = anterior cingulate cortex, PL = prelimbic, IL = infralimbic.

### Maternal bumetanide treatment prevents the deficits in behavioral flexibility seen with ethanol exposure in utero

To determine if the observed normalizing effect of maternal bumetanide treatment on tangential migration also mitigates deficits in PFC-dependent behavioral flexibility, we compared the performance of offspring exposed *in utero* to binge-type ethanol without and with maternal bumetanide treatment (0.15 mg/kg i.p, dissolved in DMSO) on the modified Barnes maze (Fig. 6). Young adult offspring exposed to binge-type ethanol *in utero* and treated maternally with bumetanide exhibited no significant difference in the number of errors committed during the training and testing phases compared with ethanol-exposed, and control, offspring (Fig. 6a; Control 14 offspring from 4 litters; EtOH 19 offspring from 4 litters; EtOH+Bumet 8 offspring from

**Figure 6.**
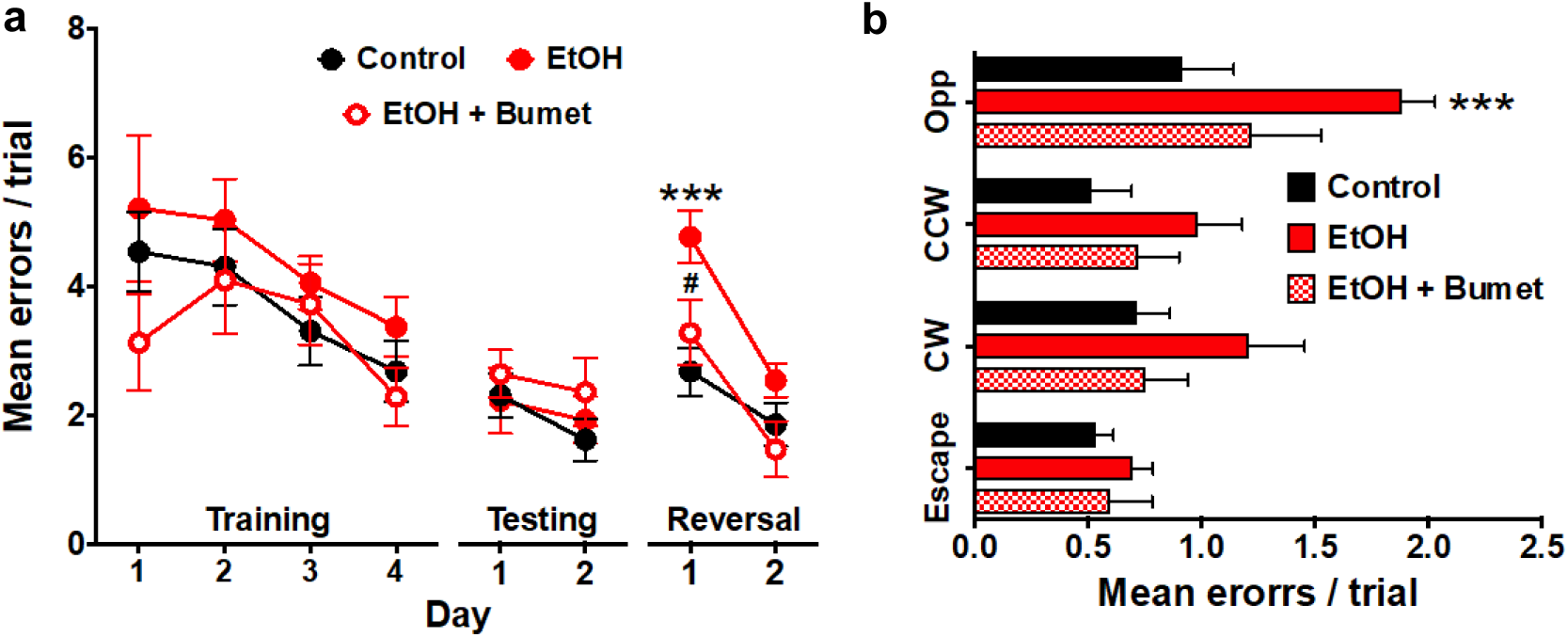
Maternal bumetanide treatment prevents the deficits in behavioral flexibility seen with ethanol exposure *in utero*. **a)** Mean number of errors committed in the modified Barnes maze by young adult mice born to control, ethanol consuming (EtOH) and ethanol consuming plus bumetanide treated (EtOH+Bumet) dams across training testing and reversal phases of testing. *** = P<0.001 compared to control, # = P<0.05 compared to EtOH; two-way ANOVA with Bonferroni post-test. **b)** Mean number of errors committed during the first reversal day stratified by quadrant relative to the escape hole. Increased errors in the opposite quadrant denotes perseverative behavior *** = P<0.001 compared to control; two-way ANOVA with Bonferroni post-test.

2 litters; P >0.427).

As previously reported (Skorput et al., 2015), the errors committed by ethanol-exposed offspring increased on day one of the reversal phase (Fig 6a; Control 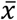= 2.68 ± 0.37 errors; EtOH 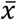= 4.78 ± 0.41 errors; P = 0.00192). Offspring born to ethanol consuming dams treated with bumetanide, however, showed no significant increase in errors compared to controls (Fig 6a; EtOH+Bumet i.p. 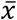= 3.28 ± 0.51 errors; P >0.999), and was significantly lower than those committed by ethanol-exposed offspring not treated maternally with bumetanide (Fig. 6b; P = 0.0433). Further analysis revealed that the increase in errors exhibited by ethanol exposed offspring on day one of the reversal phase were committed in the opposite quadrant (Fig 6b; Control 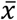= 0.92 ± 0.23 errors; EtOH 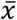= 1.88 ± 0.15 errors; P = 0.00036). In contrast, the number of these perseverative errors committed by ethanol exposed offspring maternally treated with bumetanide was not significantly different from controls (Fig. 6b; EtOH+Bumet = 1.21 ± 0.31 errors; P = 0.0745). Thus, maternal bumetanide treatment prevented perseverative behaviors, which are indicative of reduced behavioral flexibility, in offspring exposed to ethanol *in utero*.

## Discussion

The results reported here advance our understanding of depolarizing GABAergic influence on tangential migration, and demonstrate that treatment of ethanol consuming dams with bumetanide can prevent persistent interneuronopathy in ethanol-exposed offspring. These results offer insights into the developmental pathophysiology of FASD, and posit therapeutic avenues for novel treatments.

As shown by our electrophysiological studies, ethanol-induced depolarizing shifts in E_GABA_ of similar magnitude without and with bumetanide pretreatment, suggesting that the depolarizing shift of E_GABA_ induced by ethanol occurs independent of NKCC1 function. Further, as NKCC1 antagonism alone significantly shifted E_GABA_ to more hyperpolarized values, it is likely that bumetanide does not directly inhibit the ethanol-induced depolarizing shift in E_GABA_, but rather normalizes it via an opposing mechanism separate from NKCC1. This normalization of [Cl^-^]_i_, and thus the chloride driving force, had the predicted effect of attenuating GABA induced depolarizing currents in migrating cortical interneurons. Previous work by Cuzon et al. (2006, 2008) investigating the cell intrinsic and extrinsic factors that contribute to GABA’s influence on tangential migration argue strongly for GABA_A_R-activated membrane depolarization influencing both prototypical migration of GABAergic interneurons and their abnormal migration induced by ethanol exposure (Cuzon et al., 2008, 2006). The importance of GABA_A_R signaling in tangential migration is consistent with work by others describing the dependence of interneuron migration on depolarization induced calcium signaling (Behar et al., 1996). Future investigations will need to address whether the ethanol-induced potentiation of depolarizing GABA responses triggers activation of voltage-gated calcium channels and/or intracellular signaling cascades to induce aberrant migration of GABAergic interneurons.

The ability of bumetanide to prevent aberrant tangential migration following *in utero* binge-type exposure to ethanol at the height of interneuron migration sheds light on the mechanistic basis of ethanol’s enhancement of tangential migration. Maternal injection with a similar dose of bumetanide has been shown to shift the reversal potential of GABA mediated currents in neonatal cortical neurons, suggesting that maternally administered bumetanide reaches the embryonic brain (Wang and Kriegstein, 2011). With the doses of bumetanide used here, we did not see a reduction in tangential migration below levels seen in control animals, suggesting the existence of a therapeutic window in which bumetanide treatment can inhibit the aberrant migration induced by ethanol exposure *in utero* while not significantly effecting endogenous migration. Despite not impacting tangential migration *in vitro*, bumetanide at 20 µM did significantly shift the E_GABA_ of migrating interneurons to more hyperpolarized values (Fig. 1e). This discrepancy may be due to the limited magnitude of the shift in E_GABA_, and/or a difference in the acute vs persistent effects of bumetanide on E_GABA_ in MGE-derived GABAergic cortical interneurons. Further mechanistic investigations are warranted regarding the interactions of ethanol and bumetanide in augmenting E_GABA_ in both acute, and chronic settings across a range of concentrations.

Our experiments testing the long-term outcome of bumetanide treatment during binge-type ethanol exposure at the height of interneuron migration offer insights into the role of interneuronopathy in the pathophysiology of FASD, and the potential for bumetanide as a preventative treatment. The prevention of *in utero* ethanol exposure-induced changes in interneuron migration and final positioning remain correlative with the observed normalization of behavioral flexibility. However, this work strongly implicates interneuronopathy in the etiology and symptomatology of FASD. It is unlikely that aberrant tangential migration is the only effect of ethanol that contributes to the imbalance of synaptic inhibition and excitation in FASD (Kroener et al., 2012; Skorput et al., 2015). However, bumetanide’s ability to prevent this migratory defect *in vivo* suggests that such treatment may prevent ethanol-induced teratogenic effects on other cell populations that occur via potentiation of GABA_A_R activation. GABAergic interneuron migration appears to be sensitive to concentrations of ethanol that do not induce the widespread neuronal loss most commonly seen in the setting of high level exposure at a developmental time equivalent to late human gestation (Farber et al., 2010). Therefore, interneuronopathy may be an underlying etiology in a significantly larger population of FASD patients than is neurodegeneration. Dams treated with bumetanide and/or ethanol for the limited duration modeled here did not exhibit significant deviation of body weight or physical condition compared to controls. However, the influence of maternal bumetanide effects on the endpoints measured here in offspring warrant future investigation.

Preliminary studies report changes in fMRI activation and improvement in executive functioning with behavioral therapy (Nash et al., 2017). However, no treatments are currently available to ameliorate the developmental etiology of prenatal alcohol exposure. The work presented here offers antagonism of NKCC1 as a mechanism by which the short-term effects of binge-type ethanol exposure are mitigated, leading to prevention of a previously demonstrated ethanol-induced cognitive deficit. Indeed, developmental bumetanide exposure has been proposed for the treatment of neonatal epilepsy, and possible adverse effects have been reviewed (Ben-Ari, 2012; Ben-Ari and Tyzio, 2011). Our work shows that bumetanide has a biological effect on ethanol-enhanced tangential migration when administered maternally via both intraperitoneal and oral routes. While the co-administration of ethanol and bumetanide employed here served to assess mechanistic questions regarding GABAergic interneuron migration and the implications of interneuronopathy in FASD, the more likely clinical scenario would be administration of bumetanide after ethanol exposure (e.g. a patient learns she is 6 weeks pregnant, and reports binge drinking within that period of time). Therefore, the next step for this work is clearly to assess bumetanide’s ability to prevent or mitigate the long-term deleterious effects of *in utero* ethanol exposure when administered at times after ethanol exposure.

While it is likely that some enhancement of tangential migration has occurred initially, it is possible that bumetanide treatment will slow migration of subsequent interneurons to the cortex. Alternatively, it may be that bumetanide treatment later in development, when cortical neurons have already reached their final location, may allow changes in synaptic integration to prevent consolidation of what would otherwise be an imbalanced circuit. Physiological maturation of PV^+^ cortical interneurons closes the critical period of synaptic plasticity during corticogenesis (van Versendaal and Levelt, 2016). This maturation is driven by depolarizing GABAergic activity, and thus, is sensitive to the activity level of NKCC1 (Deidda et al., 2015; van Versendaal and Levelt, 2016). Indeed, work in the visual cortex demonstrates the ability for bumetanide treatment to exert lasting effects on intracortical circuit formation by extending the critical period of synaptic plasticity via a delay in the physiological maturation of PV^+^ interneurons (Deidda et al., 2015). NKCC1 antagonist treatment may therefore be beneficial for mitigating interneuronopathy directly via a delay of MGE-derived interneurons, and /or indirectly by mitigating the deleterious effects of interneuronopathy on consolidation of intracortical circuits.

## Materials and Methods

### Animals

All procedures were performed in accordance with the National Institutes of Health *Guide for the Care and Use of Laboratory Animals* and approved by the Dartmouth Institutional Animal Care and Use Committee. The Nkx2.1-Cre transgenic mouse line (originally obtained from Dr. Stewart Anderson; (Xu et al., 2008) was crossed with the Ai14 Cre-reporter mouse line (Jackson laboratories, Bar Harbor, ME) to yield Nkx2.1Cre/Ai14 mice harboring tdTomato-fluorescent (Nkx2.1^+^) MGE-derived GABAergic interneurons (Skorput et al., 2015). For time-pregnant mating, pairs of male and female mice were housed overnight, with the following day designated as E0.5, and the day of birth designated as postnatal day (P) 0. Embryonic day 13.5-16.5 embryos and P58-85 young adult mice of either sex were included in this study as available. No difference in the ratio of male to female offspring was noted between treatment groups. Embryonic day 13.5 - 16.5 and P58 - 85 were operationally defined to be within the age range equivalent to mid-first trimester (Clancy et al., 2001) and young adulthood (Varlinskaya and Spear, 2004), respectively in humans.

*Perforated patch clamp recording and drug application in acute embryonic telencephalic slices Acute slice preparation*:

On E14.5–E16.5, dams were asphyxiated with CO_2_ and fetuses were removed by caesarian section. Nkx2.1Cre/Ai14 embryos were phenotyped by the presence of tdTomato fluorescence over the cortical region of the dissected brains, which is readily visualized using UV goggles. The brains expressing tdTomato fluorescence were isolated and immersed in ice-cold oxygenated artificial cerebral spinal fluid (aCSF) containing (in mM): NaCl 124; KCl 5.0; MgCl_2_ 2.0; CaCl_2_ 2.0; glycine 0.01; NaH_2_PO_4_ 1.25; NaHCO_3_ 26; glucose 10. The brains were then embedded in 3.5% low-melting point agarose (Invitrogen, Carlsbad, CA), and coronal telencephalic slices (250 µm) were sectioned on a vibratome (Electron Microscopy Services, Hatfield, PA). The slices were stored immersed in a reservoir of aCSF at room temperature for approximately one hour prior to use for electrophysiological experiments. For consistency, we used only slices in which the MGE and LGE were clearly demarcated by the ganglionic sulcus.

### Perforated patch clamp recording

Gramicidin perforated patch clamp recording was used in order to preserve the intracellular chloride (Cl^-^) concentration (Ebihara et al., 1995). An acute 200µm telencephalic slice obtained from E14.5-E16.5 Nkx2.1Cre/Ai14 brain was transferred to a recording chamber, stabilized by an overlaying platinum ring strung with plastic threads, and maintained at 32-34° C on a heated stage fit onto a fixed-stage upright microscope (BX51WI, Olympus, Melville, NY). The slices were perfused at a rate of 0.5-1.0 ml/min with oxygenated aCSF containing (in mM): 125 NaCl, 2.5 KCl, 1 MgCl_2_, 1.25 NaH_2_PO_4_, 2 CaCl_2_·2H_2_O, 25 NaHCO_3_, 25 D-glucose, pH 7.4(adjusted with 1N NaOH). Under fluorescence illumination and Hoffman Modulation Optics (Modulation Optics, Greenvale, NY), tdTomato fluorescent cells (Nkx2.1^+^) were identified using a 40X water immersion objective (3 mm working distance; Olympus; Fig. 1a). Real-time images were captured using an analog video camera attached to a video frame grabber (Integral Technologies, Indianapolis, IN) and displayed on a computer monitor, which also aided the navigation and placement of the recording and drug pipettes.

Perforated patch-clamp recording pipettes were pulled from borosilicate glass capillaries (1.5 mm outer diameter, 0.86 mm inner diameter; Sutter Instrument Co., Novato, CA). They were first front-loaded with a K-gluconate-based internal solution containing (in mM): 100 K-gluconate, 2 MgCl_2_, 1 CaCl_2_, 11 EGTA, 10 HEPES, 30 KCl, 3 Mg^+2^ ATP, 3 Na^+^ GTP (adjusted to pH 7.3 with 1N KOH) and then back-filled with the same solution supplemented with 10μg/ml gramicidin. When filled with recording solution, the patch pipettes had resistances of 8-10 MΩ. Series resistance, monitored periodically throughout the recording, typically dropped within 10 min and stabilized for a sufficient length of time (∼15 min) for the perforated patch recording experiments. The stability of the zero current baseline was also monitored continuously, and cells with unstable recordings were excluded from analysis. Recordings were made using an AxoPatch 700B amplifier (Molecular Devices Inc., Sunnyvale, CA). Membrane currents were digitized at 20 kHz (Digidata 1320A; Molecular Devices), recorded with low-pass filtering at 10 kHz (Digidata1320A; Molecular Devices Inc.) and analyzed offline using GraphPad Prism software (Version 7.0).

### Drug application

GABA (MilliporeSigma, Burlington, MA) was dissolved in aCSF, stored as frozen stock and diluted to a working concentration of 50 μM with aCSF immediately prior to each recording session. Ethanol was prepared fresh by diluting 95% ethanol with aCSF to 6.5 mM. We used this low concentration of ethanol for the electrophysiological studies because it approximates the blood alcohol concentration (30 mg/dL) attained in a mouse model of a moderate to low level of chronic ethanol consumption used previously to investigate cortical development and function following prenatal ethanol exposure (Cuzon et al., 2008; Skorput and Yeh, 2016). Bumetanide (MilliporeSigma) was made into a salt to increase its solubility in water. A solution of sodium hydroxide was added to a suspension of bumetanide, the resultant suspension was heated to 70-80^0^C with stirring until a clear homogenate solution was obtained, which was then concentrated and dried to yield a white solid bumetanide sodium salt (3-butylamino-4-phenoxy-5-sulfamoyl-benzoic acid + Na; courtesy of Dr. Alex Pletnev, Department of Chemistry, Dartmouth College) that increased the solubility of bumetanide in water to 9 mg/ml. This bumetanide salt (MW = 362.42g/mol) was dissolved either in aCSF for the experiments involving perforated patch recording (20µM) or in liquid food in experiments that called for bumetanide being administered via the oral route to pregnant dams.

Drug solutions were loaded into separate barrels of a 6-barrel drug pipette assembly that was navigated to within 10 μm of the soma of the cell under study and applied using regulated pulses of pressure (≤ 3 p.s.i.; Picospritzer, General Valve Corporation, Fairfield, NJ) (Fig. 1a). The timing and the duration of the pressure pulses were controlled by a digital multi-channel timing unit and pulse generator (Pulsemaster A300, WPI, Sarasota, FL). One of the barrels of the multi-barrel assembly was routinely filled with aCSF, which was applied between drug applications to clear drugs from the vicinity of the cell and to control for mechanical artifacts that can occur occasionally due to bulk flow.

### Organotypic embryonic slice cultures

Time-pregnant female mice were sacrificed by carbon dioxide asphyxiation at E14.5. Embryos were removed, and the brains were harvested and processed as described previously (Cuzon et al., 2008). Briefly, the embryos were decapitated, the brains were isolated with aid of a dissecting microscope, and immersed in ice-cold slicing medium (1:1 F12:DMEM) oxygenated by bubbling with 95% O_2_ 5% CO_2_. Brains were then embedded in 3.5% low melting point agarose in 1:1 F12:DMEM. Using a vibrating microtome (Electron Microscopy Sciences), coronal slices (200µm) were collected into ice-cold slicing media saturated with 95% O_2_/5% CO_2_.

The embryonic organotypic slices were maintained on a fine nylon mesh supported on a mid-gauged U-shaped platinum wire in a 35-mm round Petri dish. A small volume of sterile filtered culture media (0.8 ml) (1:1 F12/DMEM, 1% penicillin/streptomycin, 1.2% 6 mg/ml glucose in DMEM, 10% fetal bovine serum, and 1% L-glutamine) was added to achieve air-liquid interface. Slice cultures were placed in a humidified incubator (37° C, 5% CO_2_) for 1hr, after which the following culture media were prepared and used to replenish sister slice cultures: (1) 50 mM EtOH; (2) 20 μM bumetanide; (3) 50 mM EtOH + 20 μM bumetanide; (4) vehicle (DMSO, 7.5×10^-4^ %). The concentration of ethanol to which the organotypic embryonic slice cultures were exposed was within the conventional range of concentrations employed in studies involving ethanol exposure *in* vitro (Larsen et al., 2016; Xiang et al., 2015)), and represents a value ∼3 times greater than the BAL of our binge model. The media were replenished in kind every 6 hours for the next 24 hours, when the slice cultures were washed in PBS and immerse-fixed overnight in 4% PFA/0.1 M PBS at 4°C. Following cryoprotection in 30% sucrose/0.1M PBS, the slice cultures were resectioned at 30 μm using a sliding microtome, collected in PBS, mounted on charged slides, counterstained with DAPI and coverslipped with FluorSave Reagent (Calbiochem, La Jolla, CA).

### Binge-type maternal ethanol consumption and in vivo bumetanide administration

In the *in vivo* experiments, ethanol exposure *in utero* was begun on E13.5 and terminated on E16.5 in order to be within the timeframe when tangential migration of MGE-derived cortical interneuron is at its peak in mouse (Anderson et al., 2001; Batista-Brito and Fishell, 2009; Gelman et al., 2009; Hladnik et al., 2014; Jiménez et al., 2002; Marín and Rubenstein, 2001; Parnavelas, 2000). Pregnant dams were individually housed and assigned to one of two groups: ethanol-fed, or control-fed. Mice were maintained under normal 12/12 hr light/dark cycle on a liquid diet (Research Diets, New Brunswick, NJ) supplemented with ethanol (5% w/w; ethanol-fed group) or an isocaloric control diet containing maltose (control-fed group); water was available *ad libitum*. The liquid food was replenished daily between 3:00 and 5:00 PM, when the amount consumed was measured and the dams weighed. Mice were maintained on their respective diets from E13.5 until E16.5, after which they were returned to standard chow. Dam blood alcohol level (BAL; 80±21 mg/dl) was assessed using an Analox Instruments GM7 series analyzer (Lunenburg, MA), with blood collected via the tail vein at 11:30 PM on E15.5. Dams subjected to our regimen of binge-type gestational ethanol consumption carried their offspring to full term, and litter size was unaffected (7.50 ± 0.62 for Control pups; 7.67 ± 0.76 for EtOH-exposed pups; unpaired t-test P>0.05 (Skorput et al., 2015)).

Bumetanide solution for intraperitoneal (i.p.) injection was prepared fresh from powder at the start of each binge epoch by adding bumetanide to normal saline at 0.015 mg/ml or by dissolving bumetanide in DMSO at 20 mg/ml stock concentration that was diluted to 0.015 mg/ml in normal saline for daily i.p. injections. Bumetanide was injected into binge ethanol-consuming dams at 10 ml/kg, yielding a dose of 0.15 mg/kg/day over the three-day binge period. Bumetanide delivery via liquid food was accomplished by dissolving the bumetanide sodium salt in the water used to make 5% ethanol-containing liquid food at; 1.3, 13, 130, 320 mg of bumetanide salt per kg of food.

### Imaging and analysis of immunofluorescence and Nkx2.1^+^ MGE-derived GABAergic interneurons in the prefrontal cortex

Time-pregnant dams were euthanized by CO_2_ asphyxiation on E16.5. The embryos were quickly removed, their brains dissected, immerse-fixed overnight at 4°C in PBS containing 4% paraformaldehyde (PFA)/0.1M phosphate buffered saline (PBS), and then cryoprotected in 30% sucrose/0.1M PBS. Cryosections (30μm) were cut with a sliding microtome, mounted on glass slides, DAPI counterstained and cover slipped with FluorSave Reagent (Calbiochem, La Jolla, CA). The embryonic prefrontal cortex (PFC) was delineated as part of the dorsomedial telencephalon based on DAPI counterstaining of the same sections used for counting and analyzing cells. For each embryonic brain, 10 consecutive sections of the embryonic PFC beginning at equivalent rostral-caudal levels were analyzed for counts of Nkx2.1^+^ cells.

Young adult mice were perfused trans-cardially with cold PBS and then with 4% PFA/0.1 M PBS. Brains were removed and immerse fixed in 4% PFA/0.1M PBS overnight at 4°C. Following cryoprotection in 30% sucrose, 30 μm coronal sections were made on a sliding microtome, and collected into PBS. Young adult tissue sections were blocked for 1 hour at room temperature in PBS containing 10% NGS and 0.05% Triton X-100. Then incubated overnight at 4° C with mouse anti-parvalbumin primary antibody (Life Sciences) at a dilution of 1:1000 in PBS containing 1% NGS and 0.01% Triton X-100. Following 3X washing in PBS, sections were incubated overnight with a 1:1000 dilution of Alexa Fluor 555 conjugated goat-anti-mouse secondary antibody (Invitrogen, Grand Island, NY) in PBS containing 1% NGS and 0.01% Triton X-100. Negative control with primary antibody omitted was routinely processed in parallel. For each young adult brain, sections were defined as containing the PFC beginning with the rostral most section in which all five layers of the PFC were visible, and extending caudally to the decussation of the corpus callosum. Within this operationally defined region of the PFC, images were captured from ten consecutive sections per animal beginning at equivalent rostral-caudal levels. Parvalbumin-immunopositive cells were manually counted per subdivision and per layer by trained experimenters blinded to the treatment conditions using Image J software.

Fluorescent images were captured digitally using a CCD camera (Hamamatsu, Hamamatsu city, Japan) fitted onto a spinning disk confocal microscope (BX61WI; Olympus, Melville, NY) and controlled by IPLab 4.0 software (BD Biosciences, San Jose, CA). Images were stitched using Fiji (image J, (Preibisch et al., 2009) to yield a full view of the region of interest. For *in vivo* experiments, counting of PV^+^ or Nkx2.1^+^ MGE-derived GABAergic interneurons in the embryonic or young adult PFC was automated using Fiji’s auto segmentation algorithm (RenyiEntropy) within the manually defined region of interest. For *in vitro* experiments, histological sections of organotypic slice cultures were imaged at 4X and the images were montaged to allow visualization of the cortex from the corticostriate junction to the dorsal apex. One hundred micron consecutive bins spanning the thickness of the cortex were organized along the ventral to dorsal extent of the cortex, with the starting point of the first bin aligned at the corticostriate juncture (Cuzon et al., 2006). The Nkx2.1^+^ MGE-derived GABAergic interneurons within these bins were quantified by trained experimenters blinded to the experimental condition using Fiji’s cell counting tool.

### Modified Barnes Maze

Young adult mice (P58-70) were tested behaviorally using a modified Barnes maze as previously reported (Koopmans et al., 2003; Skorput et al., 2015). Briefly, the maze consisted of twelve equally-spaced holes around a circular wall (d = 95cm), with spatial cues (large red letters) placed between the holes along the interior surface of the circular wall. Each mouse was randomly assigned an escape hole within a quadrant. All other holes were plugged. Mice were trained to find the escape hole with four 4-min trials per day for four consecutive days. Each trial ended when the mouse entered the escape hole. The mouse was then returned to its home cage (escape reinforcement). Following the training phase, the mice were rested for two days, and then tested for their ability to recall the position of the escape hole on the next two days, with four 4-min trials per day (testing phase). After two days of testing, the location of the escape hole was switched to the hole directly opposite its initial position (reversal phase). Mice were then tested for their ability to find the reversed escape hole with four 4-min trials per day over the subsequent two days.

At the beginning of each trial, the mouse was placed in the center of the maze and covered with a start box; time began when the box was removed. Latency to enter escape hole, time per quadrant; Home (H), Clockwise (CW), Opposite (Opp.), Counter Clockwise (CCW) (clock directions are relative to escape quadrant), and distance per quadrant measurements were made via video tracking software running an overhead camera hung at a set distance from the surface of the maze. During a trial, errors made (nose pokes in non-escape hole) and their locations, were recorded manually by observers on opposite sides of the arena.

### Statistics

All young adult histological data were acquired from the defined subregions and layers within the PFC. All groups consisted of data acquired from ten 30 μm tissue sections per animal (n=1: 1 animal =10 sections) from a minimum three individual animals of multiple litters. For electrophysiological data, n refers to the number of litters used in order to minimize litter effects, and the total number of cells recorded for each experiment is noted in results. Group means were compared by unpaired t-test, one-way ANOVA or two-way ANOVA with appropriate post-hoc test as indicated, and reported in figure legends, and reported as mean (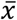) ± standard error in the results section. Reported exact P values are multiplicity adjusted for ANOVA testing of multiple comparisons. We note that previously published counts of prenatal and postnatal cortical interneurons and behavioral data obtained from control, and prenatal binge ethanol-exposed young adult mice (Skorput et al., 2015) were included for comparison with the bumetanide-treated cohort in the present study. This was justified as, in practice, the experiments involving all three cohorts were conducted in parallel, and reporting of the bumetanide data required the subsequent mechanistic studies reported here.

## Acknowledgements

The authors acknowledge funding from the National Institute on Alcohol Abuse and Alcoholism at the National Institutes of Health PHS NIH R01 AA-023410 and R21 A-024036 to H.H.Y.

## Author Contributions

**A.G.J.S**. contributed to the conception and design of the work, acquisition, analysis, and interpretation of the data, and wrote the manuscript. **S.M.L**. contributed to the design of the work, acquisition, analysis, and interpretation of the data; and writing of the manuscript.

**P.W.L.Y.** contributed to the acquisition and analysis of the data. **H.H.Y.** contributed to the conception and design of the work, interpretation of the data, and writing of the manuscript.

## References

Anderson, S.A., Marín, O., Horn, C., Jennings, K., Rubenstein, J.L., 2001. Distinct cortical migrations from the medial and lateral ganglionic eminences. Dev. Camb. Engl. 128, 353–363.

Ayoola, A.B., Stommel, M., Nettleman, M.D., 2009. Late recognition of pregnancy as a predictor of adverse birth outcomes. Am. J. Obstet. Gynecol. 201, 156.e1–156.e6. https://doi.org/10.1016/j.ajog.2009.05.011

Batista-Brito, R., Fishell, G., 2009. The developmental integration of cortical interneurons into a functional network. Curr. Top. Dev. Biol. 87, 81–118. https://doi.org/10.1016/S0070-2153(09)01203-4

Behar, T.N., Li, Y.X., Tran, H.T., Ma, W., Dunlap, V., Scott, C., Barker, J.L., 1996. GABA stimulates chemotaxis and chemokinesis of embryonic cortical neurons via calcium-dependent mechanisms. J. Neurosci. Off. J. Soc. Neurosci. 16, 1808–1818.

Ben-Ari, Y., 2014. The GABA excitatory/inhibitory developmental sequence: a personal journey.Neuroscience 279, 187–219. https://doi.org/10.1016/j.neuroscience.2014.08.001

Ben-Ari, Y., 2012. Blocking seizures with the diuretic bumetanide: promises and pitfalls. Epilepsia 53, 394–396. https://doi.org/10.1111/j.1528-1167.2011.03378.x

Ben-Ari, Y., 2002. Excitatory actions of gaba during development: the nature of the nurture. Nat. Rev. Neurosci. 3, 728–739. https://doi.org/10.1038/nrn920

Ben-Ari, Y., Khalilov, I., Kahle, K.T., Cherubini, E., 2012. The GABA excitatory/inhibitory shift in brain maturation and neurological disorders. Neurosci. Rev. J. Bringing Neurobiol. Neurol. Psychiatry 18, 467–486. https://doi.org/10.1177/1073858412438697

Ben-Ari, Y., Tyzio, R., 2011. Is It Safe to Use a Diuretic to Treat Seizures Early in Development ? Epilepsy Curr. 11, 192–195. https://doi.org/10.5698/1535-7511-11.6.192

Catterall, W.A., 2018. Dravet Syndrome: A Sodium Channel Interneuronopathy. Curr. Opin. Physiol. 2, 42–50. https://doi.org/10.1016/j.cophys.2017.12.007

Clancy, B., Darlington, R.B., Finlay, B.L., 2001. Translating developmental time across mammalian species. Neuroscience 105, 7–17.

Cuzon Carlson, V.C., Yeh, H.H., 2011. GABAA receptor subunit profiles of tangentially migrating neurons derived from the medial ganglionic eminence. Cereb. Cortex N. Y. N 1991 21, 1792–1802. https://doi.org/10.1093/cercor/bhq247

Cuzon, V.C., Yeh, P.W., Cheng, Q., Yeh, H.H., 2006. Ambient GABA promotes cortical entry of tangentially migrating cells derived from the medial ganglionic eminence. Cereb. Cortex N. Y. N 1991 16, 1377–1388. https://doi.org/10.1093/cercor/bhj084

Cuzon, V.C., Yeh, P.W.L., Yanagawa, Y., Obata, K., Yeh, H.H., 2008. Ethanol Consumption during Early Pregnancy Alters the Disposition of Tangentially Migrating GABAergic Interneurons in the Fetal Cortex. J. Neurosci. 28, 1854–1864. https://doi.org/10.1523/JNEUROSCI.5110-07.2008

Deidda, G., Allegra, M., Cerri, C., Naskar, S., Bony, G., Zunino, G., Bozzi, Y., Caleo, M., Cancedda, L., 2015. Early depolarizing GABA controls critical period plasticity in the rat visual cortex. Nat. Neurosci. 18, 87–96. https://doi.org/10.1038/nn.3890

Denny, C.H., Acero, C.S., Naimi, T.S., Kim, S.Y., 2019. Consumption of Alcohol Beverages and Binge Drinking Among Pregnant Women Aged 18-44 Years - United States, 2015-2017. MMWR Morb. Mortal. Wkly. Rep. 68, 365–368. https://doi.org/10.15585/mmwr.mm6816a1

Ebihara, S., Shirato, K., Harata, N., Akaike, N., 1995. Gramicidin-perforated patch recording: GABA response in mammalian neurones with intact intracellular chloride. J. Physiol. 484 (Pt 1), 77–86. https://doi.org/10.1113/jphysiol.1995.sp020649

Farber, N.B., Creeley, C.E., Olney, J.W., 2010. Alcohol-induced neuroapoptosis in the fetal macaque brain. Neurobiol. Dis. 40, 200–206. https://doi.org/10.1016/j.nbd.2010.05.025

Gelman, D.M., Martini, F.J., Nóbrega-Pereira, S., Pierani, A., Kessaris, N., Marín, O., 2009. The embryonic preoptic area is a novel source of cortical GABAergic interneurons. J. Neurosci. Off. J. Soc. Neurosci. 29, 9380–9389. https://doi.org/10.1523/JNEUROSCI.0604-09.2009

Hladnik, A., Džaja, D., Darmopil, S., Jovanov-Milošević, N., Petanjek, Z., 2014. Spatio-temporal extension in site of origin for cortical calretinin neurons in primates. Front. Neuroanat. 8. https://doi.org/10.3389/fnana.2014.00050

Ismail, S., Buckley, S., Budacki, R., Jabbar, A., Gallicano, G.I., 2010. Screening, diagnosing and prevention of fetal alcohol syndrome: is this syndrome treatable? Dev. Neurosci. 32, 91–100. https://doi.org/10.1159/000313339

Jiménez, D., López-Mascaraque, L.M., Valverde, F., De Carlos, J.A., 2002. Tangential migration in neocortical development. Dev. Biol. 244, 155–169. https://doi.org/10.1006/dbio.2002.0586

Kato, M., Dobyns, W.B., 2005. X-linked lissencephaly with abnormal genitalia as a tangential migration disorder causing intractable epilepsy: proposal for a new term, “interneuronopathy.” J. Child Neurol. 20, 392–397. https://doi.org/10.1177/08830738050200042001

Katsarou, A.-M., Moshé, S.L., Galanopoulou, A.S., 2017. Interneuronopathies and their role in early life epilepsies and neurodevelopmental disorders. Epilepsia Open 2, 284–306. https://doi.org/10.1002/epi4.12062

Kodituwakku, P.W., 2010. A neurodevelopmental framework for the development of interventions for children with fetal alcohol spectrum disorders. Alcohol Fayettev. N 44, 717–728. https://doi.org/10.1016/j.alcohol.2009.10.009

Koopmans, G., Blokland, A., van Nieuwenhuijzen, P., Prickaerts, J., 2003. Assessment of spatial learning abilities of mice in a new circular maze. Physiol. Behav. 79, 683–693.

Kroener, S., Mulholland, P.J., New, N.N., Gass, J.T., Becker, H.C., Chandler, L.J., 2012. Chronic alcohol exposure alters behavioral and synaptic plasticity of the rodent prefrontal cortex. PloS One 7, e37541. https://doi.org/10.1371/journal.pone.0037541

Lange, S., Probst, C., Gmel, G., Rehm, J., Burd, L., Popova, S., 2017. Global Prevalence of Fetal Alcohol Spectrum Disorder Among Children and Youth. JAMA Pediatr. 171, 948–956. https://doi.org/10.1001/jamapediatrics.2017.1919

Larsen, Z.H., Chander, P., Joyner, J.A., Floruta, C.M., Demeter, T.L., Weick, J.P., 2016. Effects of ethanol on cellular composition and network excitability of human pluripotent stem cell-derived neurons. Alcohol. Clin. Exp. Res. 40, 2339–2350. https://doi.org/10.1111/acer.13218

Marín, O., Rubenstein, J.L., 2001. A long, remarkable journey: tangential migration in the telencephalon. Nat. Rev. Neurosci. 2, 780–790. https://doi.org/10.1038/35097509

May, P.A., Baete, A., Russo, J., Elliott, A.J., Blankenship, J., Kalberg, W.O., Buckley, D., Brooks, M., Hasken, J., Abdul-Rahman, O., Adam, M.P., Robinson, L.K., Manning, M., Hoyme, H.E., 2014. Prevalence and characteristics of fetal alcohol spectrum disorders. Pediatrics 134, 855–866. https://doi.org/10.1542/peds.2013-3319

May, P.A., Blankenship, J., Marais, A.-S., Gossage, J.P., Kalberg, W.O., Joubert, B., Cloete, M., Barnard, R., De Vries, M., Hasken, J., Robinson, L.K., Adnams, C.M., Buckley, D., Manning, M., Parry, C.D.H., Hoyme, H.E., Tabachnick, B., Seedat, S., 2013. Maternal alcohol consumption producing fetal alcohol spectrum disorders (FASD): quantity, frequency, and timing of drinking. Drug Alcohol Depend. 133, 502–512. https://doi.org/10.1016/j.drugalcdep.2013.07.013

May, P.A., Chambers, C.D., Kalberg, W.O., Zellner, J., Feldman, H., Buckley, D., Kopald, D., Hasken, J.M., Xu, R., Honerkamp-Smith, G., Taras, H., Manning, M.A., Robinson, L.K., Adam, M.P., Abdul-Rahman, O., Vaux, K., Jewett, T., Elliott, A.J., Kable, J.A., Akshoomoff, N., Falk, D., Arroyo, J.A., Hereld, D., Riley, E.P., Charness, M.E., Coles, C.D., Warren, K.R., Jones, K.L., Hoyme, H.E., 2018. Prevalence of Fetal Alcohol Spectrum Disorders in 4 US Communities. JAMA 319, 474–482. https://doi.org/10.1001/jama.2017.21896

Nash, K., Stevens, S., Clairman, H., Rovet, J., 2017. Preliminary Findings that a Targeted Intervention Leads to Altered Brain Function in Children with Fetal Alcohol Spectrum Disorder. Brain Sci. 8. https://doi.org/10.3390/brainsci8010007

Owens, D.F., Kriegstein, A.R., 2002. Is there more to GABA than synaptic inhibition? Nat. Rev. Neurosci. 3, 715–727. https://doi.org/10.1038/nrn919

Parnavelas, J.G., 2000. The origin and migration of cortical neurones: new vistas. Trends Neurosci. 23, 126–131.

Peadon, E., Rhys-Jones, B., Bower, C., Elliott, E.J., 2009. Systematic review of interventions for children with Fetal Alcohol Spectrum Disorders. BMC Pediatr. 9, 35. https://doi.org/10.1186/1471-2431-9-35

Preibisch, S., Saalfeld, S., Tomancak, P., 2009. Globally optimal stitching of tiled 3D microscopic image acquisitions. Bioinformatics 25, 1463–1465. https://doi.org/10.1093/bioinformatics/btp184

Pruett, D., Waterman, E.H., Caughey, A.B., 2013. Fetal alcohol exposure: consequences, diagnosis, and treatment. Obstet. Gynecol. Surv. 68, 62–69. https://doi.org/10.1097/OGX.0b013e31827f238f

Rowles, B.M., Findling, R.L., 2010. Review of pharmacotherapy options for the treatment of attention-deficit/hyperactivity disorder (ADHD) and ADHD-like symptoms in children and adolescents with developmental disorders. Dev. Disabil. Res. Rev. 16, 273–282. https://doi.org/10.1002/ddrr.120

Skorput, A.G.J., Gupta, V.P., Yeh, P.W.L., Yeh, H.H., 2015. Persistent Interneuronopathy in the Prefrontal Cortex of Young Adult Offspring Exposed to Ethanol In Utero. J. Neurosci. Off. J. Soc. Neurosci. 35, 10977–10988. https://doi.org/10.1523/JNEUROSCI.1462-15.2015

Skorput, A.G.J., Yeh, H.H., 2016. Chronic Gestational Exposure to Ethanol Leads to Enduring Aberrances in Cortical Form and Function in the Medial Prefrontal Cortex. Alcohol. Clin. Exp. Res. 40, 1479–1488. https://doi.org/10.1111/acer.13107

van Versendaal, D., Levelt, C.N., 2016. Inhibitory interneurons in visual cortical plasticity. Cell. Mol. Life Sci. 73, 3677–3691. https://doi.org/10.1007/s00018-016-2264-4

Varlinskaya, E.I., Spear, L.P., 2004. Changes in sensitivity to ethanol-induced social facilitation and social inhibition from early to late adolescence. Ann. N. Y. Acad. Sci. 1021, 459–461. https://doi.org/10.1196/annals.1308.064

Wang, D.D., Kriegstein, A.R., 2011. Blocking Early GABA Depolarization with Bumetanide Results in Permanent Alterations in Cortical Circuits and Sensorimotor Gating Deficits. Cereb. Cortex N. Y. NY 21, 574–587. https://doi.org/10.1093/cercor/bhq124

Wang, D.D., Kriegstein, A.R., 2009. Defining the role of GABA in cortical development. J. Physiol. 587, 1873–1879. https://doi.org/10.1113/jphysiol.2008.167635

Xiang, Y., Kim, K.-Y., Gelernter, J., Park, I.-H., Zhang, H., 2015. Ethanol Upregulates NMDA Receptor Subunit Gene Expression in Human Embryonic Stem Cell-Derived Cortical Neurons. PLOS ONE 10, e0134907. https://doi.org/10.1371/journal.pone.0134907

Xu, Q., Tam, M., Anderson, S.A., 2008. Fate mapping Nkx2.1-lineage cells in the mouse telencephalon. J. Comp. Neurol. 506, 16–29. https://doi.org/10.1002/cne.21529

